# ALDH3A2 acts as a metabolic safeguard that regulates sphingolipid metabolism to suppress DNA damage and cell death

**DOI:** 10.64898/2026.07.31.741968

**Authors:** Tomoya Hotani, Maiko Sasano, Taro Okada, Taketoshi Kajimoto, Masakazu Shinohara, Satoshi Ninagawa, Tetsushi Iwasaki, Masayuki Yokoi, Kaoru Sugasawa, Wataru Sakai

## Abstract

Highly reactive aldehydes are generated during metabolic processes in the body, and their detoxification is essential for maintaining cellular homeostasis. Hexadecenal, a long-chain fatty aldehyde, is formed during the sphingolipid degradation pathway from the lipid mediator sphingosine-1-phosphate (S1P). However, the cytotoxicity resulting from dysregulation of hexadecenal metabolism is still unclear. To elucidate the effects of impaired hexadecenal metabolism, we analyzed the function of ALDH3A2, an aldehyde dehydrogenase in humans. Our results revealed that ALDH3A2 enzymatic activity is crucial for the suppression of DNA damage, particularly interstrand DNA crosslinks, upon S1P exposure. Furthermore, we demonstrated that hexadecenal accumulation promotes cell death accompanied by the activation of cellular stress responses and morphological abnormalities in the endoplasmic reticulum. These findings suggest that ALDH3A2 functions as a metabolic safeguard to suppress DNA damage and cell death in response to the enhanced metabolic flux of hexadecenal.

## Introduction

Metabolic reactions produce a variety of chemical compounds that play important roles in the body. However, some of them possess high levels of cytotoxicity if not appropriately metabolized [1]. Aldehydes are among the most common metabolites that can react with essential biomolecules such as DNA and proteins, consequently disrupting normal cellular functions. Various metabolic enzymes mitigate the detrimental damage of aldehydes. Nineteen aldehyde dehydrogenases (ALDHs) have been found in the human genome that directly catabolize aldehydes into carboxylic acids using NAD^+^ as a cofactor [2,3]. Among these, ALDH2 is the most extensively studied ALDH [4]. It oxidizes acetaldehyde produced during alcohol metabolism. *ADH5* encodes glutathione-dependent formaldehyde dehydrogenase, which belongs to the alcohol dehydrogenase family. Recent studies have shown that dysregulation of endogenous aldehyde detoxification significantly impacts human health. Mutations in both *ALDH2* and *ADH5* cause abnormalities in endogenous formaldehyde metabolism, resulting in a hereditary disorder characterized by aplastic anemia, intellectual disability, and dwarfism [5,6]. Dysregulation of endogenous formaldehyde induces DNA damage such as DNA interstrand crosslinks (ICLs), which interfere with the replication and transcription of DNA, thus adversely affecting human health. The inability to repair ICLs is linked to the pathogenesis of Fanconi anemia, which is characterized by bone marrow failure, congenital defects, and an increased risk of certain cancers [7]. Based on research from the past decades, two-tier protection against aldehyde toxicity has been proposed: enzymatic detoxification as the first tier, which metabolizes harmful aldehydes, and DNA repair mechanisms as the second tier, which repair DNA damage caused by residual aldehydes [8]. Consistent with this model, individuals with both Fanconi anemia and defective ALDH2 show accelerated progression of diseases [9].

Sjögren–Larsson syndrome (SLS) is another genetic disease caused by impaired aldehyde metabolism. It is characterized by congenital ichthyosis, spasticity, and intellectual disability due to a deficiency of fatty aldehyde dehydrogenase, ALDH3A2 [10,11]. ALDH3A2 is primarily localized in the endoplasmic reticulum (ER), with a minor splicing variant in the peroxisome. It is involved in the metabolism of long-chain fatty aldehydes. Deficiency of ALDH3A2 leads to abnormal lipid metabolism and significantly affects the skin and nervous system [12]. Hexadecenal, a major aldehyde responsible for the pathogenesis of SLS [13], is produced from sphingosine-1-phosphate (S1P) by an S1P lyase, SGPL1, in the sphingolipid degradation pathway [14]. Although the neurological abnormalities in SLS are thought to be caused by the formation of hexadecenal adducts with protein [15,16], the pathogenesis underlying its diverse clinical symptoms remains unclear. Furthermore, *in vitro* analyses revealed that α,β-unsaturated aldehydes, including hexadecenal, can form DNA adducts and ICLs [17,18]. However, it is unclear whether such DNA damage can also occur *in vivo*.

In this study, we found that ALDH3A2 deficiency led to the accumulation of hexadecenal and ICLs in cells. This dysregulation induces a characteristic type of cell death accompanied by morphological abnormalities in the ER. The results highlight the importance of the sphingolipid degradation pathway for genome stability and suppression of cell death, and propose novel insights into the sources of endogenous ICLs and the pathogenesis of SLS.

## Results

### ALDH3A2 is essential for cell survival in response to sphingolipids and hexadecenal exposure

Hexadecenal is derived from S1P in the sphingolipid degradation pathway (Figure 1A) and is one of the major substrates of ALDH3A2. Metabolic abnormalities in hexadecenal are considered to be involved in the pathogenesis of SLS. In this study, to analyze the impact of ALDH3A2 deficiency on genomic instability, ALDH3A2 knockout (ALDH3A2^KO^) cells were generated from human osteosarcoma U2OS cells that harbor wild-type p53 and represent physiological responses to DNA-damaging agents [19]. The rodent ALDH3A2^KO^ cells are sensitive to sphingolipids [16]. To determine whether human ALDH3A2^KO^ cells also exhibit sensitivity to sphingolipids, we examined their viability after continuous S1P exposure. All ALDH3A2^KO^ cells showed hypersensitivity to S1P compared to the parental U2OS cells (Figure S1A; clone #1 was used for all subsequent analyses). Furthermore, trypan blue staining showed that ALDH3A2^KO^ cells exhibit significantly increased cell death compared to the parental U2OS cells when exposed to S1P at concentrations of 5 *µ*M or higher (Figure 1B). These results indicate that the enhanced sensitivity to S1P in ALDH3A2^KO^ cells resulted from increased S1P-induced cell death. In addition to S1P, ALDH3A2^KO^ cells exhibited high sensitivity to sphingosine and hexadecenal (Figure 1C and 1D). The ectopic expression of the wild-type ALDH3A2 rescued the levels of sensitive cells. However, the expression of the catalytic-dead ALDH3A2 mutant (C241S) could not rescue cellular sensitivity, and the cells with this mutation exhibited a sensitivity comparable to that of ALDH3A2^KO^ cells. The C241S mutation completely abolishes the enzymatic activity of ALDH3A2 [20]. In contrast, ALDH3A2^KO^ cells showed no sensitivity to a small aldehyde, formaldehyde (Figure S1B). These results indicate that ALDH3A2 activity is critical in the sphingolipid degradation pathway and loss of its activity increases cellular susceptibility to sphingolipids and hexadecenal exposure.

**Figure 1.**
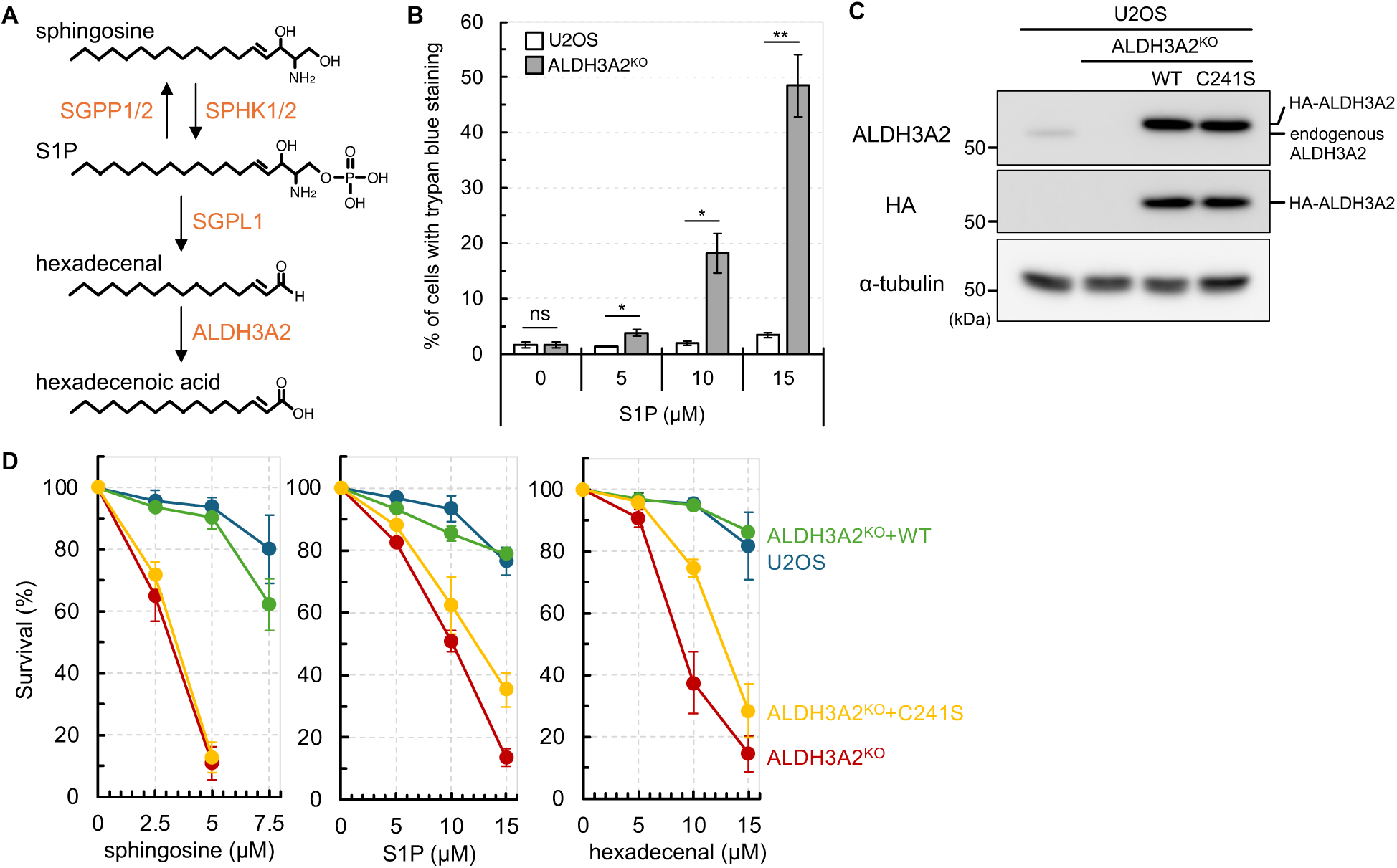
ALDH3A2 is required for cell survival after sphingolipids and hexadecenal exposure. (**A**) Schematic diagram of the sphingolipid degradation pathway. S1P can be recycled to sphingosine by S1P phosphatase 1 and 2 (SGPP1 and SGPP2) or irreversibly metabolized by S1P lyase 1 (SGPL1). The resulting fatty aldehyde, hexadecenal, is oxidized to the corresponding fatty acid by long-chain fatty aldehyde dehydrogenase, ALDH3A2. (**B**) Cell death was assessed by trypan blue staining. Cells were treated with the indicated S1P doses. After 2 days of exposure, all cells were collected and counted after trypan blue staining. Data represent mean ± standard error of the mean (SEM) from three independent experiments. Statistical significance of the differences was assessed using Student’s t-test (two-tailed). ns = not significant, \**P* < 0.05, \*\**P* < 0.01 (**C**) Immunoblot analyses of ALDH3A2^KO^ and ectopically expressed HA-tagged wild-type ALDH3A2 (WT) or mutated ALDH3A2 (C241S). α-tubulin was used as a loading control. (**D**) Cell viability was assessed after 48 h of exposure to the indicated doses (sphingosine, S1P, and hexadecenal) by crystal violet staining. Data represent mean ± SEM from at least three independent experiments.

### Accumulation of hexadecenal in ALDH3A2^KO^ cells is induced by S1P exposure

Although it has been shown that ALDH3A2^KO^ cells are susceptible to S1P exposure, the intracellular level of hexadecenal remains unclear. Therefore, using mass spectrometry, we analyzed the intracellular levels of sphingosine, S1P, and hexadecenal after S1P exposure (Figure 2). The accumulation of sphingosine and S1P was detected in an S1P-exposure-dependent manner with a higher impact on ALDH3A2^KO^ cells. However, the accumulation of hexadecenal after S1P exposure was detected only in ALDH3A2^KO^ cells. These results revealed that ALDH3A2 deficiency does not affect the conversion from S1P to sphingosine but impairs the catabolism of hexadecenal.

**Figure 2.**
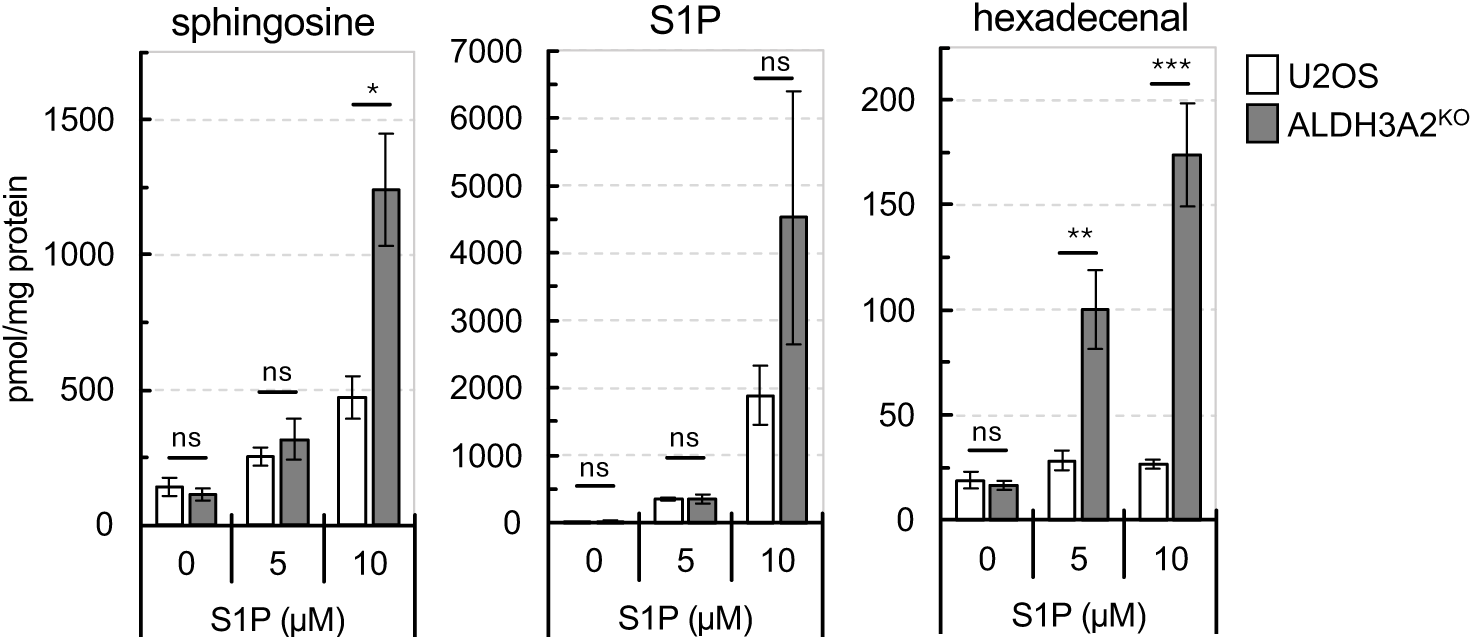
Evaluation of cellular sphingolipids and hexadecenal levels in ALDH3A2^KO^ cells after S1P exposure. Cellular sphingolipids and hexadecenal levels were analyzed using liquid chromatography-tandem mass spectrometry (LC-MS/MS). After 48 h of exposure to the indicated S1P doses, lipids were extracted from each cell line (U2OS and ALDH3A2^KO^). Data represent mean ± SEM from four independent experiments. Statistical significance of the differences was assessed using Student’s t-test (two-tailed). ns = not significant, \**P* < 0.05, \*\**P* < 0.01, \*\*\**P* < 0.001.

### ALDH3A2 deficiency promotes increased DNA damage during S1P upregulation

Hexadecenal accumulates in the ALDH3A2^KO^ cells upon exposure to S1P. Although *in vitro* studies have reported that hexadecenal can cause DNA damage [17,18], it remains unclear whether the accumulation of endogenous hexadecenal can cause DNA damage in the cells. To address this issue, we analyzed whether S1P exposure increases DNA damage using the phosphorylated histone variant H2AX (γH2AX) as a marker of DNA damage. Upon S1P exposure, the parental U2OS cells showed marked induction of γH2AX foci formation, and this effect was more enhanced in ALDH3A2^KO^ cells (Figure 3A and 3D). However, no S1P-induced γH2AX foci formation was detected in ALDH3A2^KO^ cells expressing wild-type ALDH3A2 (Figure 3B). A significant induction of γH2AX foci formation similar to ALDH3A2^KO^ cells was detected in ALDH3A2^KO^ cells expressing a mutant ALDH3A2 (C241S) lacking catalytic activity. These results indicate that S1P exposure causes DNA damage and that the enzymatic activity of ALDH3A2 is essential for its suppression.

**Figure 3.**
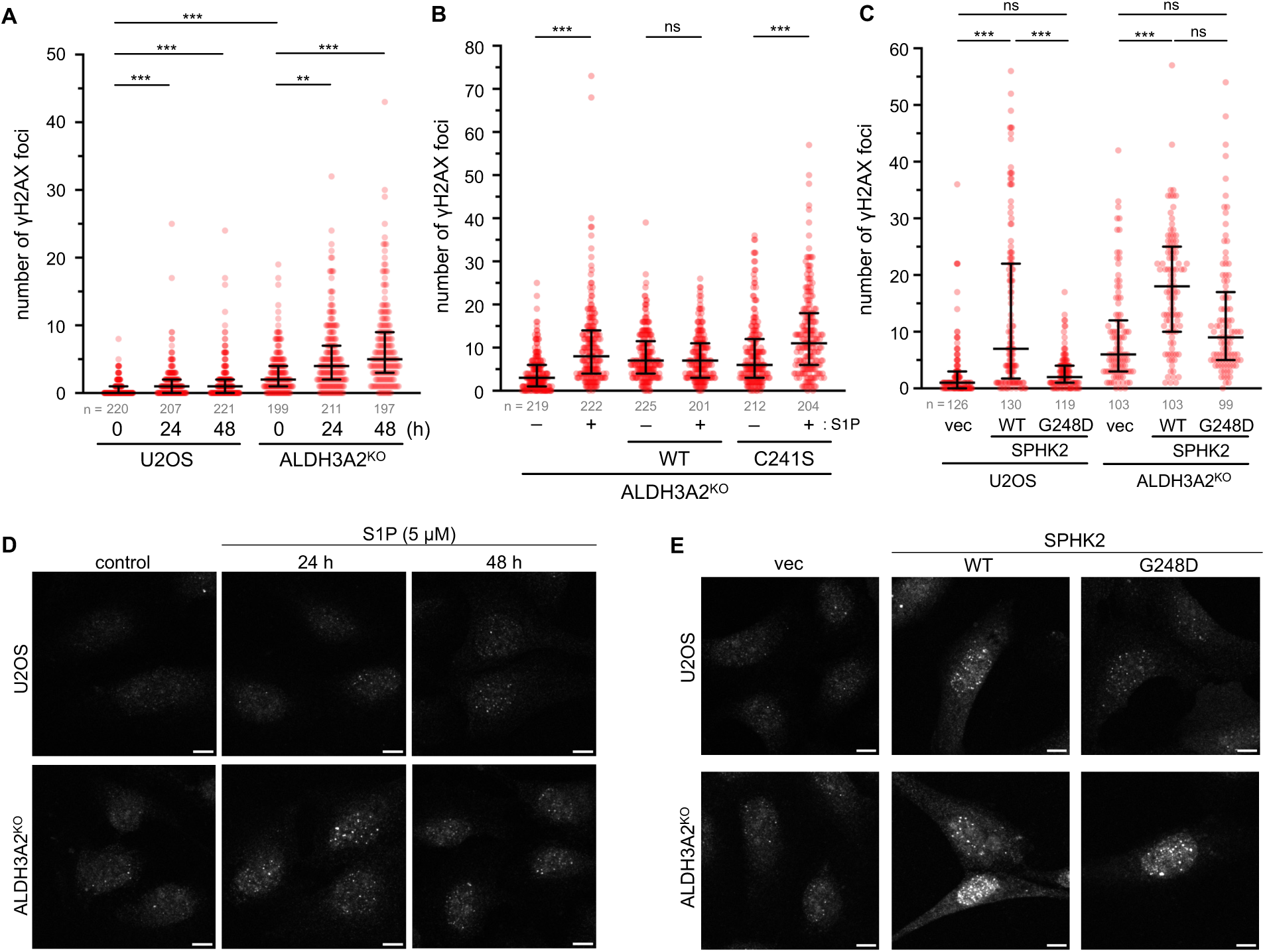
H2AX phosphorylation in response to S1P exposure and ectopic SPHK2 expression. (**A**) Quantification of S1P-induced γH2AX foci formation in U2OS and ALDH3A2^KO^ cells. Cells were treated with 5 *µ*M of S1P for the indicated time and immunostained with anti-γH2AX antibody. (**B**) Quantification of γH2AX foci formation in ALDH3A2^KO^ cells complemented with wild-type ALDH3A2 (WT) or mutant ALDH3A2 (C241S). After 48 h of exposure to S1P (5 *µ*M), cells were immunostained with anti-γH2AX antibody. (**C**) Quantification of γH2AX foci formation in U2OS and ALDH3A2^KO^ cells following transient expression of wild-type SPHK2 (WT) or kinase-inactive mutant SPHK2 (G248D). vec indicates cells transfected with an empty pEGFP vector. (**A**–**C**) Each dot represents the number of γH2AX nuclear foci per nucleus. Data are represented as median ± interquartile range. Statistical significance of the differences was assessed using the Kruskal–Wallis test followed by Dunn’s post-test for multiple comparisons. ns = not significant, \*\**P* < 0.01, \*\*\**P* < 0.001. (**D**) Representative images from (**A**) are shown. Scale bars: 10 *µ*m. (**E**) Representative images are shown in (**C**). Scale bars: 10 *µ*m.

S1P is produced from sphingosine by two sphingosine kinases, SPHK1 and SPHK2, in cells (Figure 1A). We hypothesized that transient expression of nuclear-localized SPHK2 [21] would also induce γH2AX foci formation as observed upon S1P exposure. Therefore, we transiently expressed wild-type SPHK2 and mutant SPHK2 (G248D) lacking catalytic activity [22]. In both parental U2OS and ALDH3A2^KO^ cells, the expression of wild-type SPHK2 caused a significant induction of γH2AX foci formation, whereas no effect was observed when the mutant SPHK2 was expressed (Figure 3C and 3E). Notably, enhanced induction of γH2AX foci formation was observed when wild-type SPHK2 was expressed in ALDH3A2^KO^ cells. These results demonstrate that DNA damage can occur without exposure to exogenous S1P.

### S1P exposure induces ICL lesions in ALDH3A2^KO^ cells

To reveal the type of DNA damage in cells induced by exposure to S1P, we examined the induction of DNA double-strand breaks in S1P-treated ALDH3A2^KO^ cells using a comet assay under neutral and alkaline conditions. The results demonstrated that S1P exposure did not affect the comet tail lengthening in both neutral and alkaline conditions (Figure 4A). However, tail DNA was significantly reduced in S1P-treated ALDH3A2^KO^ cells compared to the untreated cells (Figure 4A, right). Suppression of tail DNA in the alkaline comet assay is typically associated with ICLs or DNA-protein crosslinks (DPCs) [23]. Although *in vitro* formation of ICLs from hexadecenal is reported, it remains unclear whether S1P exposure induces ICL lesions in ALDH3A2^KO^ cells. Therefore, we performed a “reverse comet assay,” leveraging the fact that ICLs and DPC lesions reduce comet tail lengthening [24]. This analysis is based on the alkaline comet assay and evaluates the presence of crosslinks by observing a reduction in tail length after co-treatment with chemical agents (Figure 4B). A significant tail lengthening was detected in the alkaline comet assay using cells treated with hydrogen peroxide (Figure 4C and 4D). However, when cells were co-treated with MMC (ICL inducer) or formaldehyde (HCHO, DPC inducer), tail lengthening was significantly reduced. Furthermore, ICL- and DPC-induced reduction in tail lengthening was distinguished by treating with a proteolytic enzyme (proteinase K, ProK) before electrophoresis [25]. ProK treatment had no effect on the reduction in tail lengthening by MMC but showed a significant recovery in that caused by formaldehyde. The reverse comet assay demonstrated that S1P exposure significantly reduced tail lengthening induced by hydrogen peroxide with no effect of proteinase K (ProK) (Figure 4E and 4F). These results suggest that S1P exposure induces ICLs rather than DPCs in ALDH3A2^KO^ cells.

**Figure 4.**
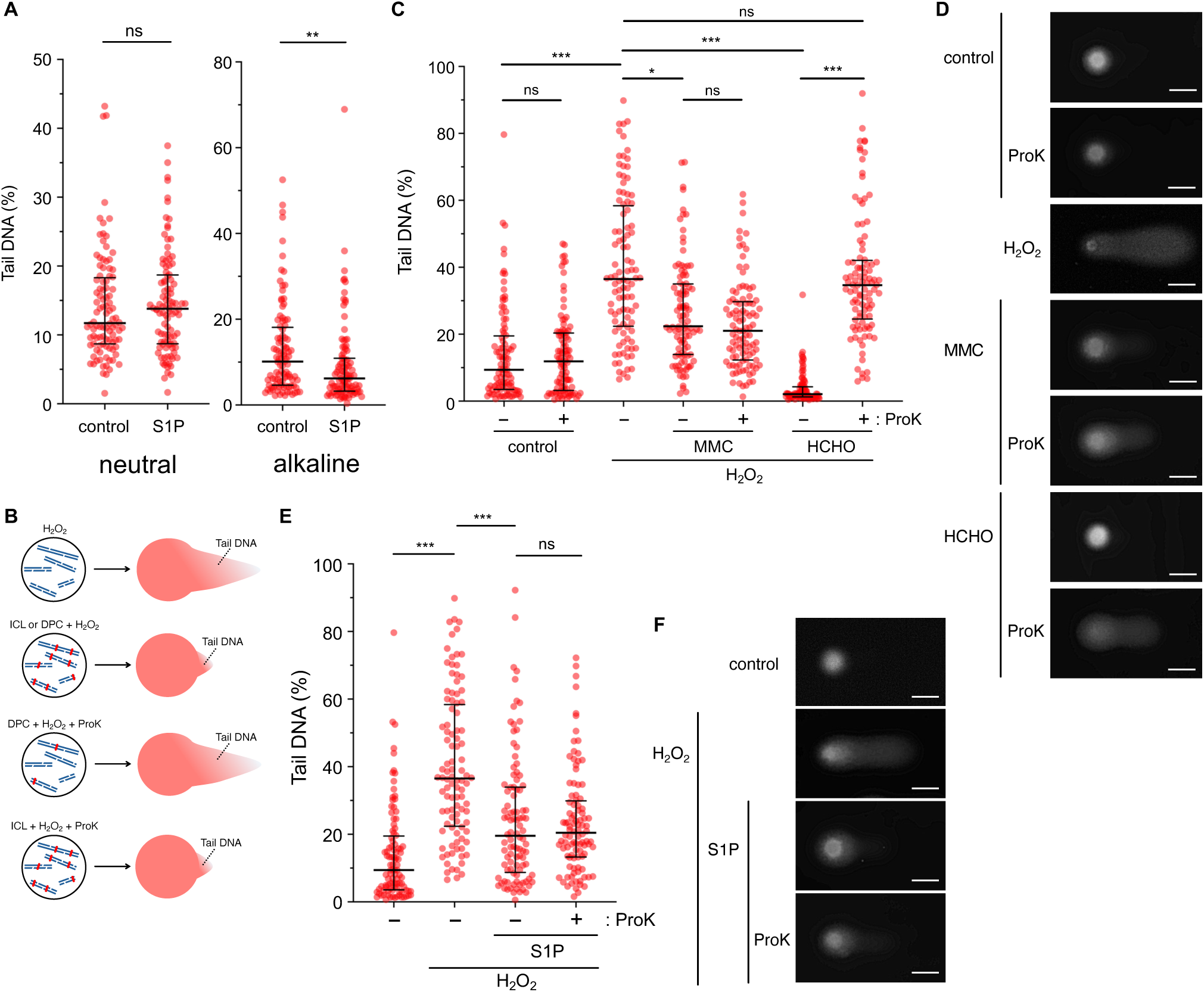
Detection of S1P-induced ICLs in ALDH3A2^KO^ cells. (**A**) Quantitation of tail DNA (%) in ALDH3A2^KO^ cells. Cells were treated with 5 *µ*M S1P for 48 h and subsequently analyzed by neutral (left) and alkaline (right) comet assays. Each dot corresponds to the tail DNA (%) of one comet. Data are represented as median ± interquartile range (n = 100). Statistical significance of the differences was assessed using the two-tailed Mann–Whitney U test. ns = not significant; \*\**P* < 0.01. (**B**) Schematic diagram of reverse comet assay principle. H_2_O_2_ exposure induced strand breaks in genomic DNA, resulting in DNA fragmentation and an increase in tail DNA under alkaline conditions. Pre-treatment with ICL or DPC-inducing agents suppressed the lengthening of tail DNA owing to the formation of crosslinking DNA fragments or proteins. Post-treatment with ProK counteracted the suppression of tail DNA lengthening by DPC crosslinking but had no effect in the condition with ICLs. (**C**) Validation of the reverse comet assay. MMC and formaldehyde (HCHO) were used as ICL and DPC-inducing agents, respectively. H_2_O_2_ exposure induced tail DNA lengthening, but co-exposure with MMC suppressed it. ProK treatment (+) had no effect on the suppression. In contrast, the suppressive effect of formaldehyde on tail DNA lengthening was counteracted after ProK treatment. The reverse comet assay with ProK treatment can distinguish DPC from ICL. Each dot corresponds to the tail DNA (%) of one comet. Data represent median ± interquartile range (n = 100). Statistical significance of the differences was assessed using the Kruskal–Wallis test followed by Dunn’s post-test for multiple comparisons. ns = not significant, \**P* < 0.05, \*\*\**P* < 0.001. (**D**) Representative comet images from (**C**) are shown. Scale bars: 50 *µ*m. (**E**) Quantification of tail DNA (%) in the reverse comet assay. After 48 h of exposure to S1P (5 *µ*M), cells were treated with H_2_O_2_ (100 *µ*M) for 1 h and analyzed by alkaline comet assay with or without ProK treatment. Each dot corresponds to the tail DNA (%) of one comet. Data represent the median ± interquartile range (n = 100). Statistical significance of the differences was assessed using the Kruskal–Wallis test followed by Dunn’s post-test for multiple comparisons. ns = not significant, \*\*\**P* < 0.001. (**F**) Representative comet images from (**E**) are shown. Scale bars: 50 *µ*m.

### S1P exposure has a marginal effect on FANCD2^KO^ cells

The Fanconi anemia pathway plays a critical role in ICL repair. FANCD2 is a key component of the Fanconi anemia pathway and forms nuclear foci in response to ICLs induced by agents such as MMC [26]. Since our results suggested S1P exposure can induce ICLs in cells, we examined the S1P sensitivity of FANCD2^KO^ cells to evaluate the contribution of the Fanconi anemia pathway to the S1P-induced ICL response. However, the effect of FANCD2 deficiency on S1P exposure was not notable (Figure 5A). Consistently, S1P exposure did not affect the FANCD2 nuclear foci formation in both parental U2OS and ALDH3A2^KO^ cells (Figure 5B). In accordance with the two-tier protection model against aldehyde cytotoxicity, FANCD2 single KO itself may not affect S1P sensitivity. Therefore, we generated FANCD2 ALDH3A2 double knockout (DKO) cells from FANCD2^KO^ cells and examined their S1P sensitivity. However, none of the DKO clones exhibited higher S1P sensitivity than the ALDH3A2^KO^ cells, nor did they show a synergistic effect (Figure 5A and Figure S1A). These results indicate that S1P exposure increases γH2AX levels but does not activate the damage response via the Fanconi anemia pathway.

**Figure 5.**
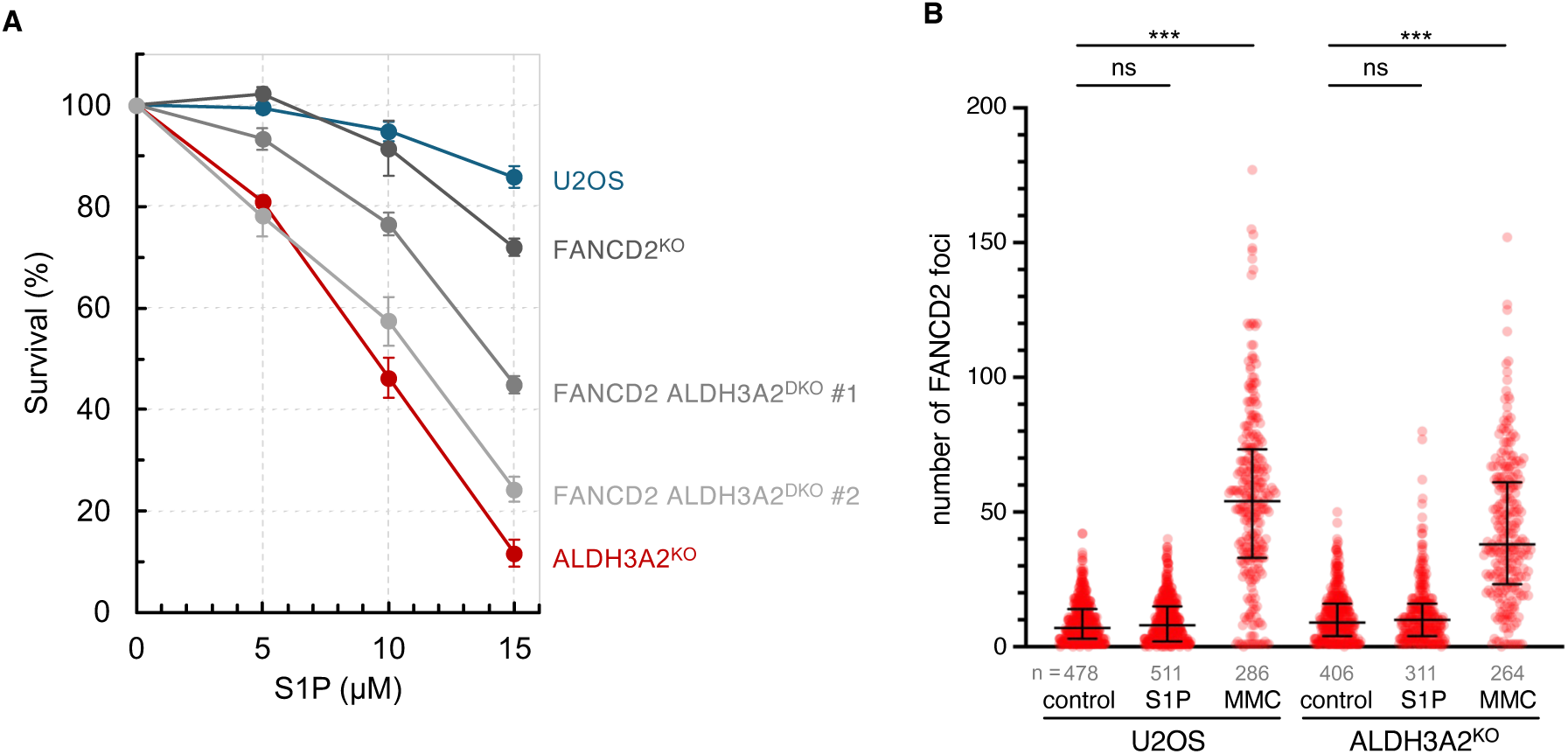
S1P exposure has a marginal effect on FANCD2^KO^ cells. (**A**) Cell viability was assessed after 48 h of exposure to the indicated S1P doses by crystal violet staining. Two FANCD2 ALDH3A2^DKO^ clones with distinct mutations were generated from FANCD2^KO^ cells. Data represent mean ± SEM from at least three independent experiments. (**B**) Quantification of S1P-induced FANCD2 foci in U2OS and ALDH3A2^KO^ cells. Cells were treated with 5 *µ*M of S1P for 2 days or 300 nM of MMC. Each dot represents the number of FANCD2 nuclear foci. Data represent the median ± interquartile range. Statistical significance of the differences was assessed using the Kruskal–Wallis test followed by Dunn’s post-test for multiple comparisons. ns = not significant, \*\*\**P* < 0.001.

### S1P-induced cell death is accompanied by morphological abnormalities in the ER

The results showed that ALDH3A2^KO^ cells are highly sensitive to S1P exposure, thereby causing cell death rather than cell-cycle arrest (Figure 1B). To investigate whether S1P exposure induced apoptosis, we assessed PARP1 cleavage by caspases as an apoptosis marker. The results revealed that S1P exposure induced PARP1 cleavage in ALDH3A2^KO^ cells but not in parental U2OS cells (Figure 6A). These results were consistent with the proportion of dead cells upon S1P exposure (Figure 1B). Next, we examined whether caspase activation is required for S1P-induced cell death using Z-VAD-FMK, a pan-caspase inhibitor. S1P-induced PARP1 cleavage was inhibited in a Z-VAD-FMK-dependent manner (Figure 6B). However, ALDH3A2^KO^ cells exhibited high S1P sensitivity even under anti-apoptotic conditions (Figure 6C, right). These results suggest that S1P exposure induces apoptosis in ALDH3A2^KO^ cells, but it is not required for cell death.

**Figure 6.**
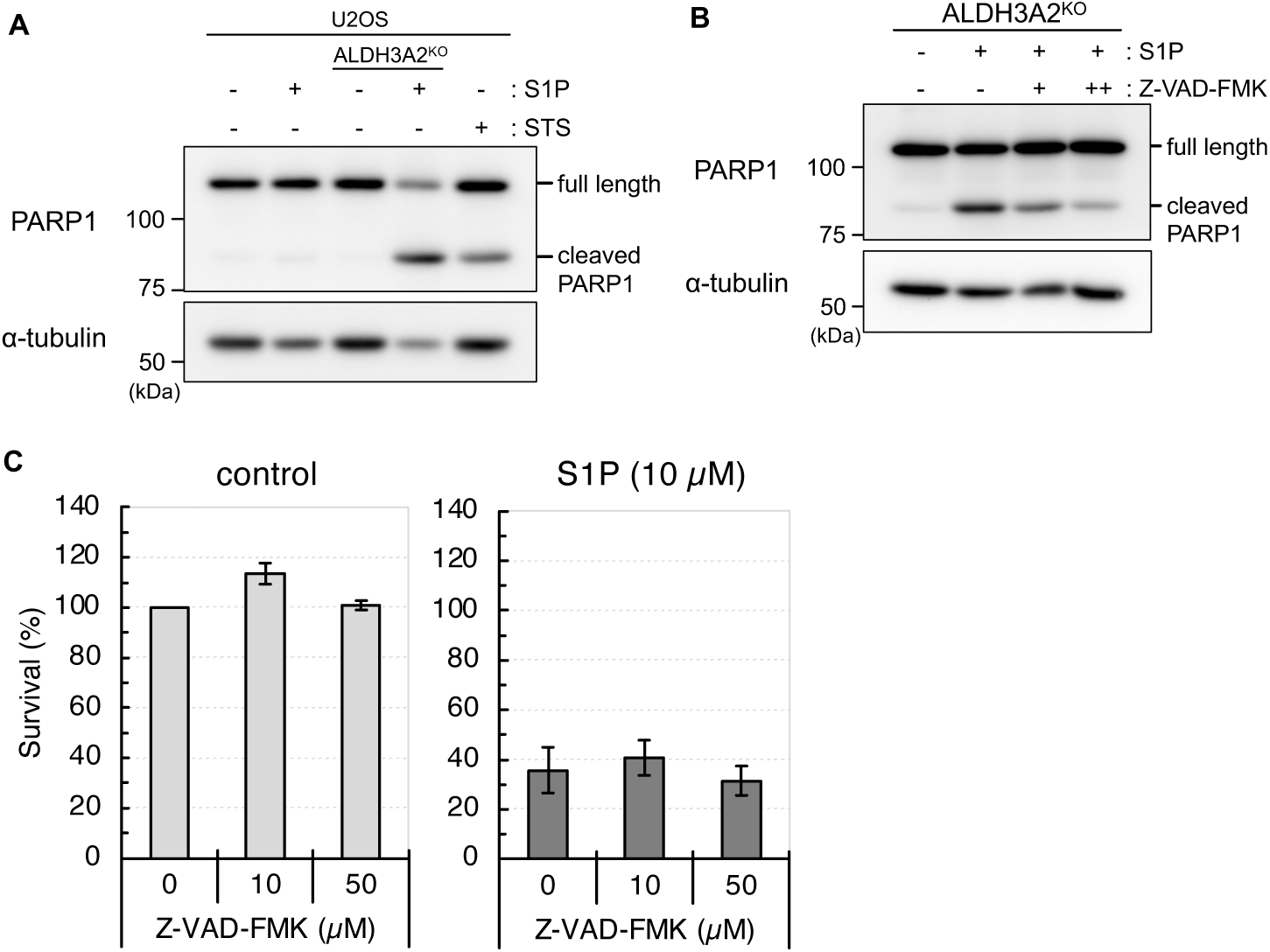
S1P-induced cell death in ALDH3A2^KO^ cells. (**A**,**B**) Cleavage of PARP1 after S1P exposure. Cell lysates were subjected to immunoblotting with the indicated antibodies. α-tubulin was used as a loading control. (**A**) The parental U2OS and ALDH3A2^KO^ cells were treated with 15 *µ*M of S1P for 24 h. Staurosporine (STS) was used as an apoptosis inducer (1 *µ*M, 6 h). (**B**) ALDH3A2^KO^ cells were treated with 15 *µ*M of S1P and Z-VAD-FMK (+: 10 *µ*M, ++: 50 *µ*M) for 24 h. Z-VAD-FMK was used as a pan-caspase inhibitor. (**C**) The effect of a caspase inhibitor on S1P-induced cell death. Cell viability was assessed after 48 h exposure with 10 *µ*M of S1P and/or indicated Z-VAD-FMK doses by crystal violet staining. Left graph (control) indicates cell viability treated with only Z-VAD-FMK. Right graph (S1P) indicates cell viability co-treated with S1P and Z-VAD-FMK. Data represent mean (% of untreated) ± SEM from at least three independent experiments.

In response to exposure with high concentrations of S1P, characteristic morphological abnormalities were observed in ALDH3A2^KO^ cells (Figure S2A). Numerous vacuole-like structures were observed in the cytoplasm of ALDH3A2^KO^ cells treated with a lethal concentration of S1P (15 *µ*M), some of which affected the morphology of the nucleus. ALDH3A2 is localized in the ER, so the effects of metabolic abnormalities in ALDH3A2^KO^ cells upon S1P exposure are expected to dominate in the ER. The vacuole-like structures observed in ALDH3A2^KO^ cells may represent morphological abnormalities in the ER. We generated ALDH3A2^KO^ cells that stably expressed DsRed-KDEL (a fluorescent protein fused to an ER localization signal) and analyzed ER localization and morphology after exposure to a lethal level of S1P. The results showed that the vacuole-like structures were swollen ERs (Figure 7A). Furthermore, ALDH3A2^KO^ cells with morphological abnormalities in the ER appeared 24 h or longer after exposure to lethal levels of S1P. The morphological abnormalities of the ER subsequently underwent rapid shrinkage (Figure 7B). Paraptosis is a type of cell death accompanied by ER swelling [27]. Mitochondrial swelling, another morphological hallmark of paraptosis, was also observed in ALDH3A2^KO^ cells upon S1P exposure (Figure 7C).

**Figure 7.**
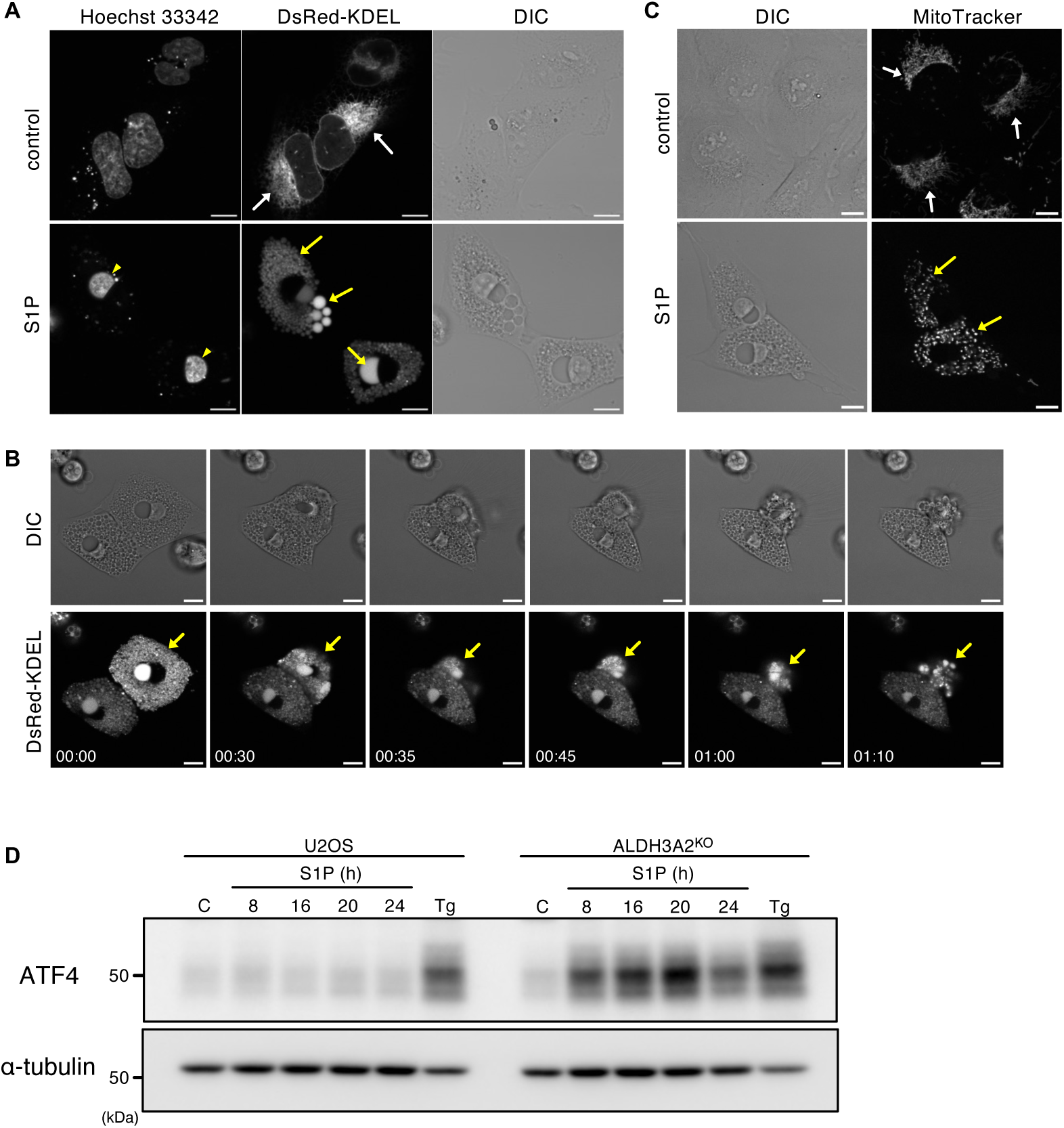
Morphological abnormality of ER and cellular stress response in response to S1P exposure in ALDH3A2^KO^ cells. (**A**) Representative images of ALDH3A2^KO^ cells expressing DsRed-KDEL as a probe for the ER lumen. ALDH3A2^KO^ cells showed normal ER morphology in the regular medium (white arrows). After 24 h exposure to S1P (15 *µ*M), a part of ALDH3A2^KO^ cells showed a small nucleus (yellow arrowheads) and dilated ER (yellow arrows). Scale bars: 10 *µ*m. (**B**) Live-cell imaging of cell shrinkage in ER-swelling ALDH3A2^KO^ cells (yellow arrows). Selected frames after 24 h of exposure to S1P (15 *µ*M) are shown. Numbers indicate time (hr:min) from the first frame. Scale bars: 10 *µ*m. (**C**) ALDH3A2^KO^ cells showed normal mitochondrial morphology in the regular medium (white arrows). After 24 h exposure to S1P (15 *µ*M), mitochondrial swelling occurred in the ER-dilated ALDH3A2^KO^ cells (yellow arrows). Scale bars: 10 *µ*m. (**D**) Induction of ATF4 expression after S1P exposure. The parental U2OS and ALDH3A2^KO^ cells were treated with 15 *µ*M of S1P for the indicated times. A control sample was prepared from untreated cells. Thapsigargin (Tg) was used as an ER stress inducer (300 nM, 8 h). Cell lysates were subjected to immunoblotting with an ATF4 antibody. α-tubulin was used as a loading control. Representative images are shown.

### S1P exposure triggers cellular stress responses in ALDH3A2^KO^ cells

To maintain cellular homeostasis in response to various endogenous and exogenous stresses, cells activate an adaptive pathway termed the integrated stress response (ISR) [28]. To verify whether the ISR is activated in ALDH3A2^KO^ cells upon S1P exposure, we examined the induction of ATF4, a main effector of the ISR. The results revealed that S1P exposure did not induce ATF4 expression in the parental U2OS cells. However, ATF4 induction was detected in ALDH3A2^KO^ cells 8 h after S1P exposure, with the highest expression observed after 20 h (Figure 7D). The ISR is activated in response to various stresses, including amino acid starvation, viral infection, heme deficiency, and ER stress. S1P exposure and hexadecenal accumulation are likely to affect a wide range of cellular processes. Based on the morphological abnormalities induced by S1P exposure and the central role of ER in S1P metabolism, we hypothesize that S1P exposure triggers ER stress. ATF6α is an ER membrane–bound stress sensor that undergoes cleavage under ER stress [29]. Therefore, we assessed whether S1P exposure induces ATF6α cleavage. The ATF6α cleavage products were detected in ALDH3A2^KO^ cells in a pattern similar to the induction of ATF4 expression (Figure S2B). Together, these findings suggest that S1P exposure induces ER stress and activates ISR before the onset of paraptosis-like cell death in ALDH3A2^KO^ cells.

## Discussion

S1P is a lipid mediator that is present ubiquitously both inside and outside the cell. Its metabolic pathways have been extensively studied. S1P is recycled in sphingolipid biosynthesis via dephosphorylation and is irreversibly degraded from S1P to hexadecenal via the degradation pathway, the only exit pathway for sphingolipid metabolism. ALDH3A2 metabolizes a long-chain fatty aldehyde, hexadecenal, into fatty acids. In this study, we demonstrated that S1P exposure induces marked accumulation of hexadecenal, resulting in enhanced cytotoxicity in ALDH3A2^KO^ cells. In addition, ALDH3A2 deficiency leads to increased DNA damage, particularly ICLs, as well as paraptosis-like cell death accompanied by ER swelling via S1P upregulation.

Aldehydes are an endogenous source of ICLs. α,β-unsaturated aldehydes, such as acrolein and 4-hydroxyl-2-nonenal, derived from lipid peroxidation, form propano-2’-deoxyguanosine adducts that can produce ICLs [18]. Here, we focused on a sphingolipid-derived long-chain fatty aldehyde, hexadecenal, which possesses an unsaturated bond at the α-β position and is known to form DNA adducts *in vitro* [17]. However, it remains unclear whether endogenous hexadecenal forms DNA adducts and/or induces ICL formation in cells. Formaldehyde has been reported as a source of endogenous DNA damage [30]. A recent study also revealed that aminoaldehyde from the polyamine metabolic pathway is a genotoxic aldehyde [31]. Our results raised the possibility that hexadecenal derived from sphingolipid metabolism is a source of endogenous ICLs. S1P is produced from sphingosine by SPHK1/2 and converted to hexadecenal by SGPL1 in the ER. SPHK1 is localized predominantly to the cytoplasm [32], while SPHK2 is localized to the nucleus in a cell cycle-dependent manner [33]. SGPL1 also functions in the nucleus [34]. Therefore, hexadecenal may be produced not only in the ER but also in the nucleus. These results demonstrated that transient overexpression of SPHK2 resulted in a significant induction of γH2AX foci formation, even in the absence of S1P exposure.

The Fanconi anemia pathway is involved in response to ICLs and their repair. FANCD2^KO^ cells showed low sensitivity to S1P exposure. In accordance with the two-tier protection model against aldehyde toxicity [8], the effects of S1P exposure (accumulation of hexadecenal) may be counteracted by the activity of ALDH3A2 in FANCD2 single KO cells. Therefore, we generated FANCD2 ALDH3A2 DKO cells; however, the DKO cells exhibited S1P sensitivity equivalent to that of ALDH3A2 single KO cells. Consistently, S1P exposure did not induce nuclear foci formation of FANCD2. Why is the Fanconi anemia pathway not activated despite increased γH2AX induction by S1P exposure? The human genome contains four ALDH3 family genes: *ALDH3A1*, *ALDH3A2*, *ALDH3B1*, and *ALDH3B2*, with *ALDH3B2* being a pseudogene [2]. These ALDH3 family members share high amino acid homology and are considered to possess functional redundancy. Therefore, ALDH3A2 deficiency may be partially compensated for by other ALDH3 family members. On the other hand, biochemical analyses suggest that acetaldehyde-derived ICLs may be unhooked via an unknown repair pathway independent of the Fanconi anemia pathway [35]. It is also possible that S1P-derived ICLs are repaired via this repair pathway. Alternatively, *in vitro* analyses have demonstrated that ICLs derived from α,β-unsaturated aldehydes are chemically unstable [18]; therefore, S1P-induced ICLs could spontaneously unhook without activating the Fanconi anemia pathway.

Sphingolipid metabolic homeostasis plays an important role in cell fate [36–39]. While the accumulation of ceramides and sphingosine induces cell death, S1P acts as a signal for cell proliferation and survival. Therefore, the balance between interconvertible sphingolipids determines whether a cell lives or dies. Thus, the regulation of sphingolipid metabolism is referred to as the “sphingolipid rheostat.” In this study, we demonstrated that hexadecenal accumulation and apoptosis are induced in ALDH3A2^KO^ cells in response to S1P exposure. These results were consistent with previous findings that SGPL1 expression and hexadecenal exposure induce apoptosis [40,41]. Inhibition of apoptosis did not affect S1P-induced cell death, indicating it contributes little to S1P-induced cell death in ALDH3A2^KO^ cells. Furthermore, S1P-induced cell death, which is accompanied by characteristic morphological abnormalities in the ER and mitochondria, is distinct from the typical morphological features of apoptosis. Instead, it corresponds to a non-apoptotic form of cell death known as paraptosis [27]. Although paraptosis is triggered by various signals, ER stress is considered one of its major initiating factors. Our results demonstrated that S1P exposure induces ER stress and triggers ISR in ALDH3A2^KO^ cells. ALDH3A2 contributes to the alleviation of ER stress associated with lipid metabolism [42], supporting the possibility that ALDH3A2 maintains ER function through sphingolipid homeostasis.

The sphingolipid rheostat describes the dynamic balance among the interconvertible ceramide, sphingosine, and S1P. Although S1P is considered a pro-survival sphingolipid, our findings suggest that increased flux through the S1P degradation pathway can promote cellular stress responses and paraptosis-like cell death when downstream ALDH3A2-dependent detoxification is impaired. In other words, ALDH3A2 acts as a “metabolic safeguard” that determines the homeostasis of sphingolipid metabolism by regulating the metabolic flux of hexadecenal.

Our findings suggest that S1P-induced hexadecenal may be a source of endogenous ICLs. Further studies are required to determine whether hexadecenal-induced ICLs can be detected in genomic DNA extracted from S1P-treated cells. Furthermore, the repair pathway responding to S1P-induced DNA damage remains largely unknown. It is necessary to identify the factors that function in response to S1P-induced DNA damage. The main clinical symptoms of SLS are ichthyosis, spasticity, and intellectual disability. The S1P-induced cellular stress responses and paraptosis-like cell death may contribute to the pathogenesis of SLS. However, the analyses were based on acute cellular responses under S1P exposure at non-physiological concentrations. Notably, the physiological concentration of S1P differs across tissues and organs. The highest concentration of S1P is found in the blood, reaching approximately 1 *µ*M [43]. In the future, an analysis of the chronic effects of physiological S1P concentrations is necessary. The mechanisms underlying S1P-induced cellular stress responses and paraptosis-like cell death are unclear. Given that some neurodegenerative diseases are caused by dysregulation of ISR and ER stress [44], the relevance of S1P-induced cellular stress responses to the pathophysiology of SLS must be elucidated.

## Materials and Methods

### Plasmids

HA-tagged wild-type ALDH3A2 and the C241S mutant were cloned into pIREShyg3 (Clontech). The ALDH3A2 (C241S) mutant was generated using the QuikChange Lightning Site-Directed Mutagenesis Kit (Agilent). To generate pSPHK2(WT)-EGFP, wild-type SPHK2 cDNA was cloned into pEGFP-N1 (Clontech) as described previously [22]. The cDNA of mutant SPHK2 (G248D) [22] was cloned into pEGFP-N1. pDsRed-KDEL was kindly provided by Tamotsu Yoshimori (Osaka University). All transfections were performed using FuGENE HD Transfection Reagent according to the manufacturer’s instructions (Promega).

### Cell culture and reagents

The human osteosarcoma cell line U2OS (ATCC, HTB-96) and its derivatives were cultured at 37°C in a humidified atmosphere containing 5% CO_2_ in Dulbecco’s modified Eagle’s medium (Shimadzu Diagnostics) supplemented with 10% fetal bovine serum (Corning). Sphingosine-1-phosphate (62570, Cayman Chemical) was dissolved in dimethyl sulfoxide (DMSO):1 M HCl (95:5, v/v). Sphingosine (10007907, Cayman Chemical), hexadecenal (17566, Cayman Chemical), MMC (139-18711, Fujifilm Wako), and thapsigargin (T9033, Merck) were dissolved in DMSO. Z-VAD-FMK was purchased from Selleck Chemicals (S7023). Staurosporine was obtained from Merck (569396).

### Gene disruption

The endogenous *ALDH3A2* gene was disrupted in the U2OS and FANCD2^KO^ cell lines using the GeneArt CRISPR Nuclease Vector with CD4 Enrichment Kit (Thermo Fisher Scientific). The guide RNA targeting exon 5 of the *ALDH3A2* gene (5′-AATATAGTCGGGTGCAATGC-3′) was designed using CHOPCHOP [45]. Following enrichment with anti-CD4 magnetic beads, single clones were isolated by limiting dilution. ALDH3A2 expression in each clone was assessed by immunoblotting, and gene disruption was confirmed using Sanger sequencing of genomic DNA. FANCD2^KO^ cells were generated from U2OS cells in a previous study [46].

### Immunoblotting

Cell lysates were prepared with NP40 lysis buffer (25 mM Tris-HCl pH 7.4, 150 mM NaCl, 1 mM EDTA, 1% NP-40, and 5% glycerol) or CSK buffer (10 mM PIPES-NaOH pH 6.8, 3 mM MgCl_2_, 1 mM EGTA, 300 mM NaCl, 10% glycerol, 0.1% Triton X-100, and 50 mM NaF) containing protease inhibitors (0.25 mM phenylmethylsulfonyl fluoride, 1 μg/mL leupeptin, 2 μg/mL aprotinin, 1 μg/mL pepstatin, and 50 μg/mL Pefabloc SC). After incubation on ice for 30 min, the cell lysates were centrifuged for 10 min at 20,000× g to obtain soluble cell extracts. Proteins were separated by sodium dodecyl sulfate (SDS)-polyacrylamide gel electrophoresis and transferred onto polyvinylidene difluoride membranes (Immobilon-P, Merck). After blocking with 5% skim milk in TBS-T (50 mM Tris–HCl pH 8.0, 150 mM NaCl, and 0.1% Tween 20), the membranes were incubated overnight at 4°C with primary antibody diluted in blocking solution. Following extensive washing, the membranes were incubated with the appropriate secondary antibodies. Immunoreactive signals were visualized by chemiluminescence using CDP-Star (Thermo Fisher Scientific) as the substrate. Chemiluminescence signal was detected using a LAS4010 imaging system (Cytiva). For representative images, the brightness was adjusted using Fiji (ImageJ) [47]. Antibodies against the following proteins were used in this study: ALDH3A2 (1:2000; ab184171, Abcam), HA (1:10000; M180-3, Medical & Biological Laboratories), α-tubulin (1:20000; T5168, Merck), PARP1 (1:2000; #9542, Cell Signaling Technology), ATF4 (1:2000, sc-200; Santa Cruz Biotechnologies), and ATF6α [29]. Alkaline phosphatase-conjugated secondary antibodies were purchased from Merck (1:20000; A3688 and A3812).

### Crystal violet assay

Cell viability was determined by crystal violet assay as described previously [48], with minor modifications. Cells were seeded into 24-well plates at 1.5 × 10^4^ cells per well. After an overnight culture, the medium was replaced with fresh medium containing each drug at the indicated concentrations. After the incubation period indicated in the corresponding figure legend, the attached cells were fixed for 5–10 min in a solution containing 10% methanol and 10% acetic acid. The fixed cells were subsequently stained with 1% crystal violet (038-04862, Fujifilm Wako) in methanol. The adsorbed dye was resolubilized in 4% ethanol containing 1% SDS. The dye was transferred to 96-well plates and analyzed photometrically at 595 nm using a microplate reader.

### Quantification of sphingolipids and hexadecenal

Sphingosine and S1P were quantified by liquid chromatography-tandem mass spectrometry (LC-MS/MS) as described previously [49], with some modifications. Briefly, cells were harvested by adding two volumes of ice-cold methanol and 10 *µ*L of internal standard mixture (Splash Lipidomics Cer/Sph Mixture II, LM6005), followed by centrifugation (15,000× g for 10 min) to precipitate proteins. The supernatant was transferred to an autoinjector vial for analysis. The system consisted of a Q-Trap 6500 (Sciex) equipped with a Shimadzu LC-30AD HPLC system. A ZORBAX Eclipse Plus C18 column (100 mm × 4.6 mm, 3.5 *µ*m, Agilent Technologies) was used for sample separation. The mobile phase consisted of (A) methanol/acetonitrile/water (1:1:3) and (B) isopropanol, both containing 5 mM ammonium acetate, 500 nM EDTA, and 0.025% NH_3_ water, with a gradient from 0% to 95% of B at a 0.4 mL/min flow rate. Hexadecenal was quantified by LC-MS/MS as described previously [50], with some modifications. Cells were harvested by adding two volumes of ice-cold methanol with 10 *µ*L of internal standard mixture (10 pg/*µ*L ethanol solution of 2E-hexadecenal-d5, 857461P, Avanti Polar Lipid), followed by centrifugation (15,000× g for 10 min) to precipitate proteins. Then, 100 *µ*L of supernatant was converted to the semicarbazone derivatives by heating with 400 *µ*L of 5 mM semicarbazide hydrochloride in methanol containing 5% formic acid at 40°C for 2 h. The derivatives were subjected directly to LC-MS/MS analysis. 2E-hexadecenal was separated on an Acquity UPLC Peptide BEH C18 column (50 × 2.1 mm; 1.7 μm; Waters, USA). The column was maintained at 45°C at a 0.3 mL/min flow rate. The mobile phases consisted of (A) 1:1:3 (v/v/v) acetonitrile:methanol:water with ammonium formate (5 mM) and 10 nM EDTA, and (B) 100% isopropanol with ammonium formate (5 mM). Mass spectrometric detection of lipids was performed using multiple reaction monitoring.

### Immunostaining and cell imaging

Cells were seeded in 35 mm glass-bottom dishes (Matsunami Glass) and incubated under the conditions described in the corresponding figure legend. Fixation and immunofluorescence staining were performed as described previously [51]. Antibodies against the following proteins were used in this study: FANCD2 (1:500; NB100-182, Novus Biologicals) and phospho-H2AX (1:500; 05-636, Merck). Alexa Fluor secondary antibodies were purchased from Thermo Fisher Scientific (1:1000; A-11020, A21206). MitoTracker Green FM (M7514, Thermo Fisher Scientific) was used for mitochondria labeling. Hoechst 33342 (H342, Dojindo Laboratories) and DAPI (D212, Dojindo Laboratories) were used for counterstaining. Single-plane images were acquired using an Olympus FV3000 confocal laser scanning microscope (Evident) with a 40× objective lens. The nuclear foci formation was quantified using Fiji (ImageJ) [47].

### Comet assay

Neutral and alkaline comet assays were performed using the CometAssay kit (Trevigen) with minor modifications. Cells were pretreated with 5 *µ*M S1P for 2 days. Subsequently, cells were resuspended in molten LMAgarose (Trevigen) and allowed to solidify on glass slides (Matsunami Glass). For reverse comet assays [23–25], cells were post-treated with 100 *µ*M of hydrogen peroxide (081-04215, Fujifilm Wako) for 1 h after S1P exposure. The treated cells were processed according to the standard protocol for the alkaline comet assay (Trevigen). Before electrophoresis, cells were treated with 1 mg/mL of proteinase K (EO0491, Thermo Fisher Scientific) for 2 h in a moist chamber at 37°C to distinguish between DPCs and ICLs. The images were captured with a BZ-9000 (Keyence). The percentage of DNA in the comet tail was calculated using the OpenComet plugin for Fiji [52]. For each condition, 100 comets were analyzed. Statistical analysis was done using GraphPad Prism 9 (GraphPad Software).

### Assessment of Cell Death

Cell death was evaluated using the trypan blue exclusion method. All cells were collected from each dish. The cell pellet was then resuspended in a 0.5% trypan blue solution (204-21102, Fujifilm Wako). Cell death was determined by calculating the proportion of dead cells relative to the total number of cells.

### Data availability

This study includes no data deposited in external repositories. The data supporting the findings of this study are available from the corresponding author upon reasonable request.

## Acknowledgments

The authors are grateful to the members of the Biosignal Research Center and the Department of Biology, Graduate School of Science, Kobe University, for helpful discussions and encouragement, and to the Core Facility Center of Kobe University for supporting the experiments in this study.

## Funding

This work was supported by JST SPRING grant number JPMJSP2148 (T.H.), JSPS KAKENHI grant numbers JP20K06487 and JP25K15454 (W.S.). and partially supported by Japan Science and Technology Agency Moonshot R&D Grant Number JPMJPS2022 (M.S.).

## Author contributions

**Tomoya Hotani**: Formal analysis; Funding acquisition; Investigation. **Maiko Sasano**: Formal analysis; Investigation. **Taro Okada**: Conceptualization; Formal analysis; Investigation; Methodology. **Taketoshi Kajimoto**: Conceptualization; Formal analysis; Investigation; Methodology. **Masakazu Shinohara**: Formal analysis; Funding acquisition; Investigation; Methodology. **Satoshi Ninagawa**: Resources. **Tetsushi Iwasaki**: Resources. **Masayuki Yokoi**: Resources; Supervision. **Kaoru Sugasawa**: Resources; Supervision. **Wataru Sakai**: Conceptualization; Data curation; Formal analysis; Funding acquisition; Investigation; Methodology; Project administration; Supervision; Visualization; Writing–original draft; Writing–review & editing

## Disclosure and competing interests statement

The authors declare no competing interests.

## Supplementary Figures

**Figure S1.**
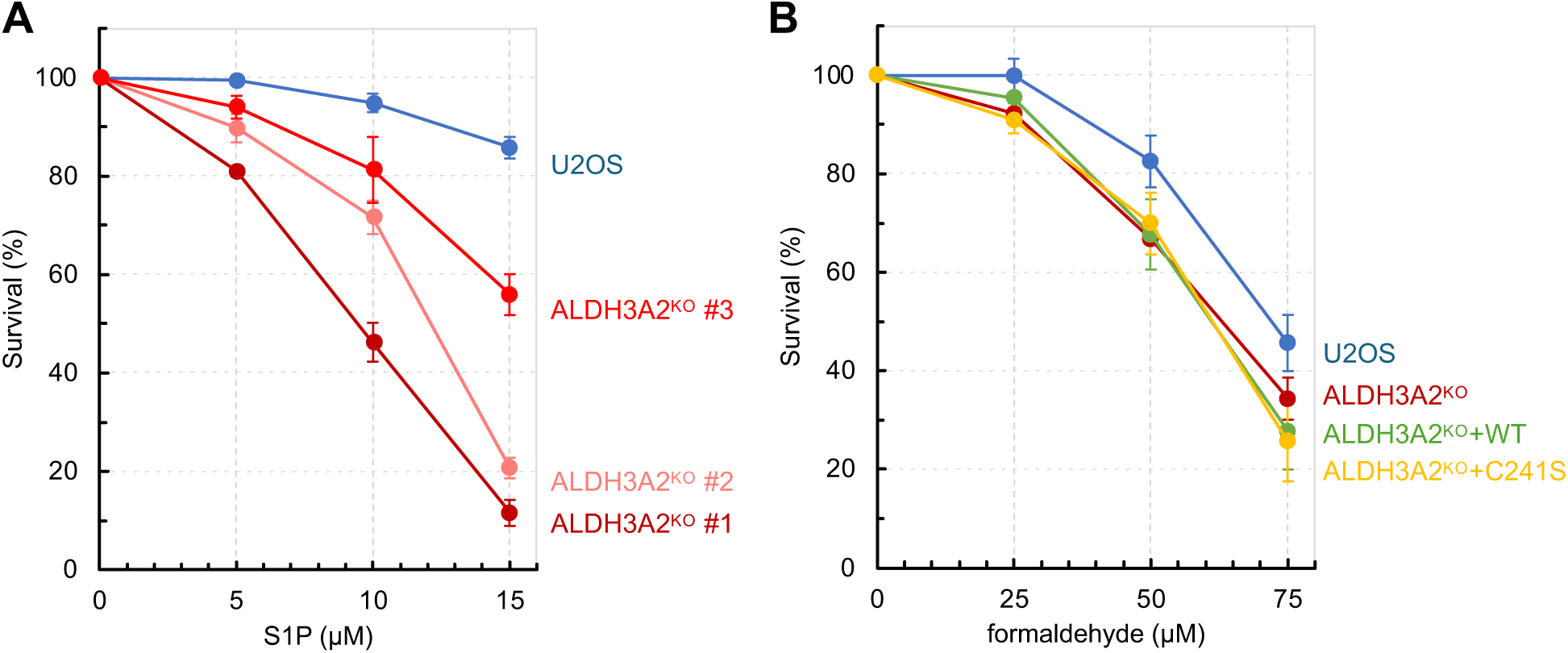
S1P sensitivity of U2OS and ALDH3A2^KO^ clones. (**A**) Cell viability was assessed after 48 h exposure to the indicated S1P doses by crystal violet staining. Three ALDH3A2^KO^ clones with distinct mutations were generated from parental U2OS cells. Clone #1 was used in this study. Data represent mean ± SEM from more than 3 independent experiments. (**B**) Cell viability was assessed after 48 h exposure to the indicated formaldehyde doses by crystal violet staining. Data represent mean ± SEM from at least three independent experiments.

**Figure S2.**
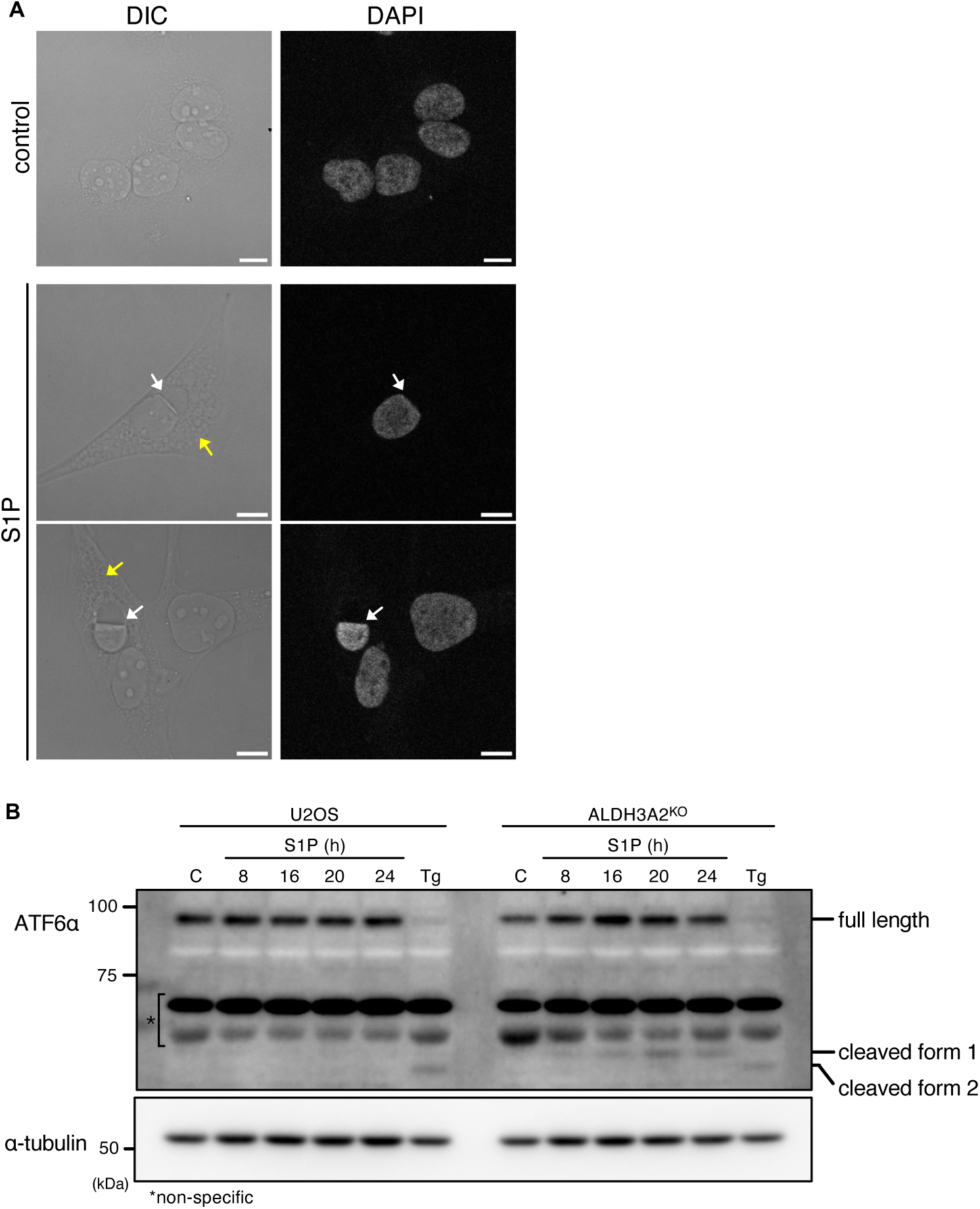
S1P-induced morphological abnormalities and ER stress response in the ALDH3A2^KO^ cells. (**A**) After 24 h exposure to S1P (15 *µ*M), ALDH3A2^KO^ cells showed atypical nuclei (white arrows) and vacuole-like structures (yellow arrows). Scale bars: 10 *µ*m. (**B**) Cleavage of ATF6α after S1P exposure. The parental U2OS and ALDH3A2^KO^ cells were treated with 15 *µ*M of S1P for the indicated times. S1P exposure induced a cleaved form 1 of ATF6α [53] in ALDH3A2^KO^ cells. The control sample was prepared from untreated cells. Tg was used as an ER stress inducer (300 nM, 1 h). Cell lysates were subjected to immunoblotting with an ATF6α antibody. α-tubulin was used as a loading control. Representative images are shown.

